# Human Tear Metabolomics Using Liquid Chromatography-Q Exactive-HF Mass Spectrometry

**DOI:** 10.1101/2021.01.18.424378

**Authors:** Gauri Shankar Shrestha, Ajay Kumar Vijay, Fiona Stapleton, Russell Pickford, Nicole Carnt

**Affiliations:** School of Optometry and Vision Science, UNSW Sydney, Australia; Bioanalytical Mass Spectrometry Facility, UNSW Sydney, Australia

**Author notes:** **Corresponding Author** Gauri S. Shrestha, M. Optom, FAAO, Scientia PhD candidate, School of Optometry and Vision Science, UNSW SYDNEY NSW 2052 AUSTRALIA. Funding: No funding was provided for the study.

**Keywords:** metabolites, tears, orbitrap, amino acids, QExactive-HF, carbohydrate, carnitine

## Abstract

**Aim:** To putatively identify and characterise human tear metabolites in a normal subject on an untargeted platform of liquid chromatography-Q exactive-HF mass spectrometry.

**Methods:** Four samples of unstimulated tears were collected from both eyes on four consecutive days between 1 – 2 pm using a microcapillary tube and pooled from both eyes each day. Untargeted analysis of the tears was performed by chromatographic separation of constituent metabolites in both CSH-C18RP (Charged Surface Hybrid-C18 Reversed Phase) and SeQuant ZIC-pHILIC (Zwitterionic-polymeric Hydrophilic Interaction Liquid Chromatography) columns, followed by heated electrospray ionization (HESI) and the acquisition of mass spectra using QExactive-HF mass spectrometer. Compound Discoverer software (v2.0) was used for data analysis.

**Result:** Eighty-two metabolites were tentatively identified. Seventy compounds (85.4 %) were observed in all four samples with a coefficient of variation (CV) less than 25 %. Fifty-nine metabolites (71.9 %) were novel in the healthy tears. Amino acids were the most frequently detected metabolites in the tears (28 %), followed by carbohydrates (12.2 %), carboxylic acids (8.5 %), carnitines (6.1 %) and glycerophospholipids (4.9 %), respectively.

**Conclusion:** The current untargeted platform is capable of detecting a range of tear metabolites across several biological categories. This study provides a baseline for further ocular surface studies.

## Introduction

Tear fluid is a complex biochemical mixture containing electrolytes, proteins, metabolites, lipids and mucins (Ablamowicz and Nichols, 2017; Chen et al., 2011; Dor et al., 2019; Rantamaki et al., 2011; Zhou et al., 2012). The tear composition reflects a relative state of ocular homoeostasis and health and can be affected by various factors such as diseases, nutrition, environment and medication (Azkargorta et al., 2017; Hagan et al., 2016; Hohenstein-Blaul et al., 2013). The tear fluid is the most easily and non-invasively accessible extracellular fluid, which makes it a suitable candidate for investigating biomarkers for diagnostic and prognostic purposes (Pieragostino et al., 2015). A biomarker is defined as “a characteristic that is objectively measured and evaluated as an indicator of normal biological processes, pathogenic processes, or pharmacologic responses to a therapeutic intervention” (Group et al., 2001).

Tear fluid has frequently been used as an easily accessible analyte pool in the study of metabolites. Around 100 tear metabolites have been reported in healthy individuals (Chen et al., 2011; ChenZhuo et al., 2000; Choy et al., 2001; Dammeier et al., 2018; Farkas et al., 2003; Gogia et al., 1998; Nakatsukasa et al., 2011; Pescosolido et al., 2009; Pintor et al., 2002; Speek et al., 1986; Van Haeringen and Glasius, 1977). Metabolites are the end products of complex cellular regulatory networks and are expressed as an influence of a combination of gene, tissue, microbiota and environment (Pinu et al., 2019). The analysis of metabolites gives insight into the physiological, pathological and biochemical status, and enhances our understanding of disease mechanism and progression (Hohenstein-Blaul et al., 2013; Pieragostino et al., 2015; Pinu et al., 2019).

The present study has profiled tear metabolites from a healthy subject collected over four consecutive days with an untargeted metabolomic platform. Untargeted metabolomics compares the relative abundance of metabolites and relative change in metabolite peak area in multiple samples without prior identification or optimisation and maximises the the number of metabolites observed compared to targeted metabolomics does (Nalbantoglu, 2019; Roberts et al., 2012).

## Methods

### Subject and tear collection

Ethics approval for the recruitment of participant was obtained from the Human Research Ethics Committee of the University of New South Wales (UNSW), Sydney (HC 180515). The study adhered to the Declaration of Helsinki. Written informed consent was obtained before enrolment into the study. Four tear samples (volume of tear > 10 μL / sample) were collected from both eyes of a male participant (age 40 years) on four consecutive days using microcapillary tubes. The participant was healthy with no ocular and systemic diseases, had no ocular trauma, surgery or injury. Every day the tear samples from both eyes were pooled. The tear sample was placed on ice and stored at −80°C until analysis.

### Chemicals

High performance liquid chromatographic grade (HPLC) acetonitrile, ammonium formate and formic acid were purchased from Sigma Aldrich (St. Louis, MO). HPLC grade methanol and isopropyl alcohol were purchased from Fisher Scientific (Pittsburgh, PA). Deionised water was collected from a cartridge-deioniser from MilliQ (Millipore; Billerica, MA).

### Metabolite extraction from tear samples

The frozen tear fluids were thawed on ice at 4°C before metabolite extraction. A 5 μL aliquot of the tear fluid was transferred into two separate tubes (sample set-1 and sample set-2) for two separate conditions of mass spectrometry. Each tube was then filled with 200 μL of 9:1 methanol: water and centrifuged for 10 minutes at 15,000 rpm, and the supernatants pipetted out immediately. The supernatants were lyophilised in a vacuum concentrator (Speed Vac, Thermo Savant, NY) at a temperature lower than 10 °C for 12 hours. Two blank samples with 200 μL of 9:1 methanol: water underwent a similar procedure to be used as controls. The resultant dried pellets from sample set-1 and sample set-2 were reconstituted in 50 µL water and 50 µL of acetonitrile, respectively. Finally, these samples were vortexed immediately before metabolites analysis.

### Liquid-chromatography-Q exactive-HF mass spectrometry (QE-HF-MS)

Chromatographic separation was performed using an Ultimate 3000 System (ThermoFisher Scientific, CA) with ACQUITY UPLC CSH-C18RP (Charged Surface Hybrid-C18 Revere Phase) Column 130 Å, 1.7 µm, 2.1 mm X 100 mm (Waters, Milford, MA) for sample set-1 and with SeQuant ZIC-pHILIC (Zwitterionic-polymeric Hydrophilic Interaction Liquid Chromatography) 5μm, 100 × 2.1 mm PEEK coated polymeric column (Merck KGaA, Darmstadt, Germany) for sample set-2.

#### Sample set-1

The LC flow rate was maintained at 0.3 ml/min. Mobile phase A was 0.1% formic acid in water and mobile phase B was 0.1% formic acid in acetonitrile, with the gradient profile of 2% B from 0 to 2 mins, 15% B at 6 mins, 50% B at 12 mins, 95% B from 16 to 18.5 mins and 2% from 19 to 25 mins.

#### Sample set-2

The LC flow rate was maintained at 0.2 ml/min. Mobile phase A was 2 mM ammonium formate in water (pH 6.6) and B, 9:1 acetonitrile: 2 mM aqueous NH_4_COOH (pH 6.6). The gradient profile was 90% B at 0 min to 80% B at 8 min, 10% B from 15 to 18 min, 90% B at 19-24 min.

Column temperature for both sample sets was maintained at 45 °C, and autosampler temperature was maintained at 4 °C. Each sample was analysed in both positive-and negative-ionisation modes, using ThermoFisher QExactive HF with HESI. The volume of sample injection was 5 μl. Each ionisation mode comprised of injection of triplicates of each sample. The mass spectrometry full scan comprised of a mass to charge ratio (m/z) between 50 and 750. Tandem mass spectrometric data (MS/MS) was acquired automatically throughout the run using the Xcalibur software’s Data Dependant Analysis. The system was set up for an automated series of LC-MS/MS samples using the ThermoFisher Xcaliber software. The sequence of the sample run was randomised using the RANDBETWEEN function in Excel.

### Metabolite detection and tentative identification

Compound Discoverer v2.0 was used for data analysis. The untargeted metabolomics workflow used is presented in Figure 1 (ThermoScientific, 2019). The workflow began with selecting input files which were acquired from an LC/MS experiment, and that contained high-resolution accurate-mass scans (MS). The multiple input files were read, filtered and aligned using the MS scan data. These steps were followed by detection and grouping of unknown compounds, compound prediction and compound annotations (name, formula, composition and structure). Tentative identification of compounds was performed using online database (ChemSpider and mzCloud) and local database (Metabolika, Mass Lists and mzVault) searches. Finally, the tentatively identified compounds were mapped to biochemical pathways using Metabolika, the Kyoto Encyclopedia of Genes and Genomes (KEGG) and the BioCyc database. The mass tolerance was set to 3 ppm for compound detection and identification.

**Figure 1.**
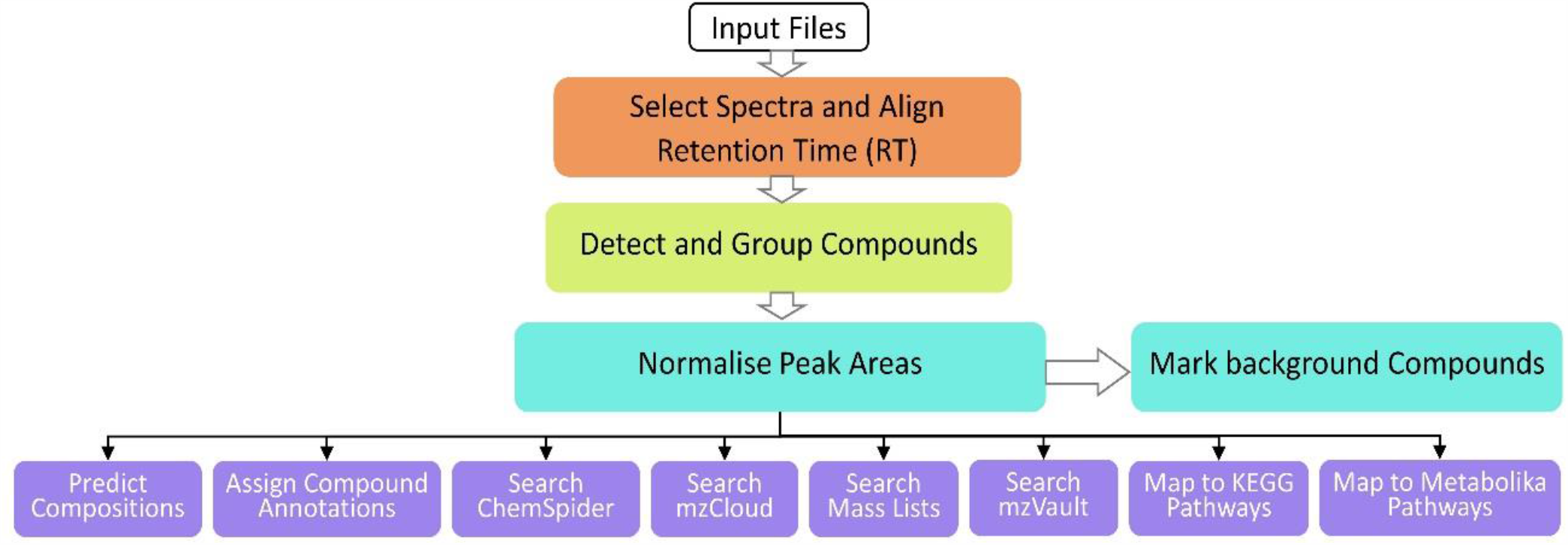
Showing workflow trees for untargeted metabolomics analysis

### Statistical Analysis

Initially, approximately 6000 candidate compounds were observed in both CSH-C18RP and ZIC-pHILIC columns under both positive and negative ionisation. After filtering out candidates with no proposed identification, around 1300 compounds remained. Further various levels of filters were applied to detect compounds in Compound Discoverer v2.0, that included a) differential analysis using ANOVA with Tukey HSD post-hoc for fold-change, the peak area of compound detection and compound ratio; b) Benjamini-Hochberg (B-H) correction to account for false discovery rate (FDR); c) compound detection in at least three samples with the CV less than or equal to 25%; and d) removing exogenous compounds such as sedatives, analgesics, herbicides, phytotoxins, and food additives. This resulted in 82 significant compounds. Principal Component Analysis was used to determine the difference between tear samples and control samples.

## Results

### Differentiation of tear samples using principal component analysis (PCA)

Before principal component analysis, compounds without proposed identity, adjusted P-value more than 0.05 and group CV more than 25 % were removed from the result lists. Data with negative fold-change for the tear samples and fold-change significant for the control sample (false-positives) were also removed. Principal component analyses obtained for CSH-C18RP and ZIC-pHILIC in positive ionisation mode for tear samples and controls were presented in figure 2, which indicated more than 50 % of the variation between samples and controls.

**Figure 2.**
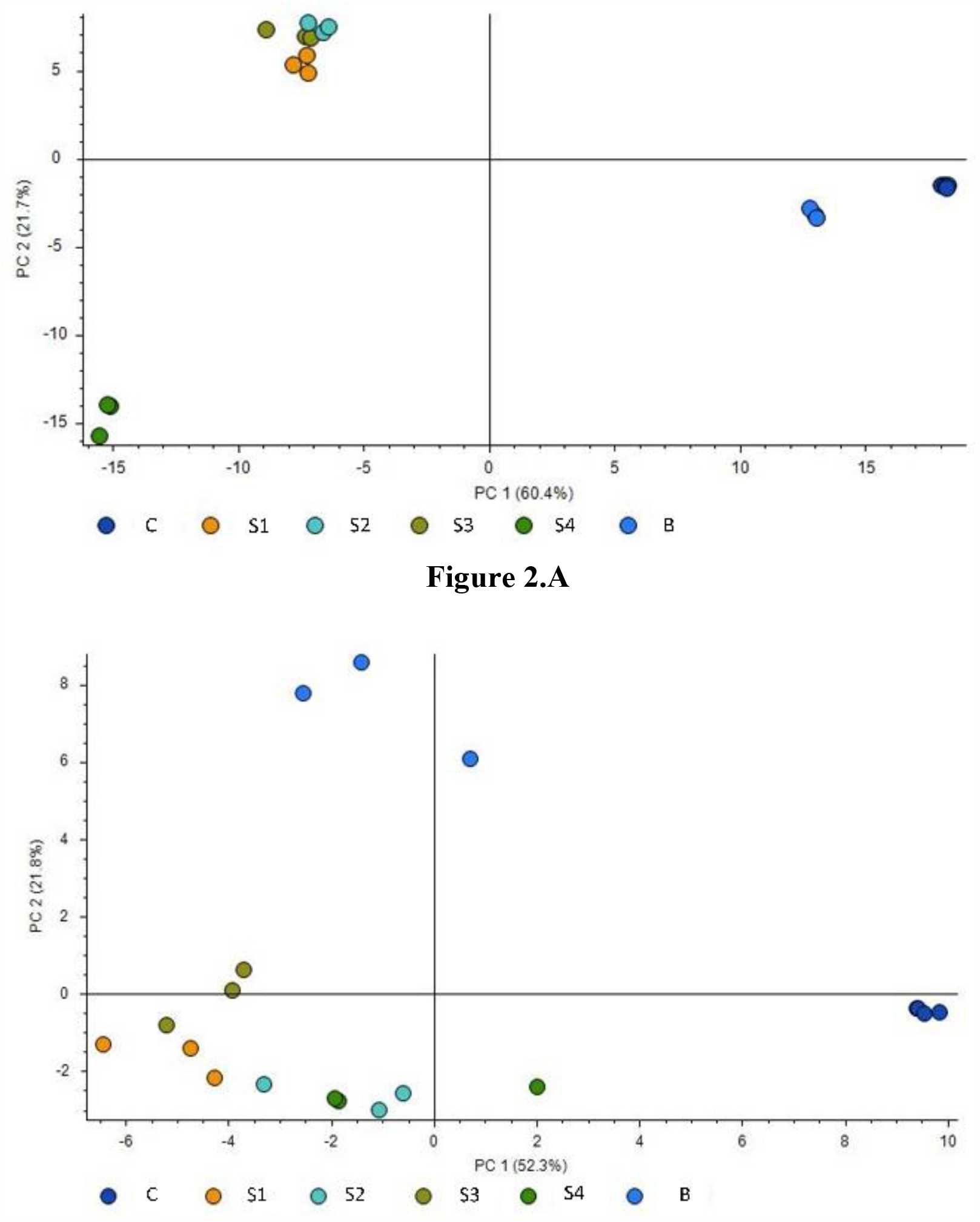
Principal component analysis showing a good separation between tear samples and a control sample for the compounds detected by mass-spectrometry in A) CSH-C18RP (+) ion mode and B) ZIC-pHILIC (+) ion mode. PC1 constitutes more than 50% of the variation between sample groups. C = control, S1-S4 = samples, B = blank (acetonitrile)

## Chromatographic separation of tear metabolites

Of 82 metabolites putatively annotated in the four ion modes. CSH-C18RP (+) and ZIC-pHILIC (+) were the most efficient in the extraction of tear metabolites (Figure 3), which resulted in sole identification of 33 and 19 metabolites, respectively. Conversely, CSH-C18RP (-) and ZIC-pHILIC (-) eluted 9 and 5 metabolites, respectively. Additionally, CSH-C18 (+) and pHILIC (+) complemented to further identification of five metabolites (N-methyl ethanolamine phosphate, 1-3-di-o-Tolylguanidine, L-pyroglutamic acid, 4-Heptyloxyphenol and L-Tyrosine).

**Figure 3.**
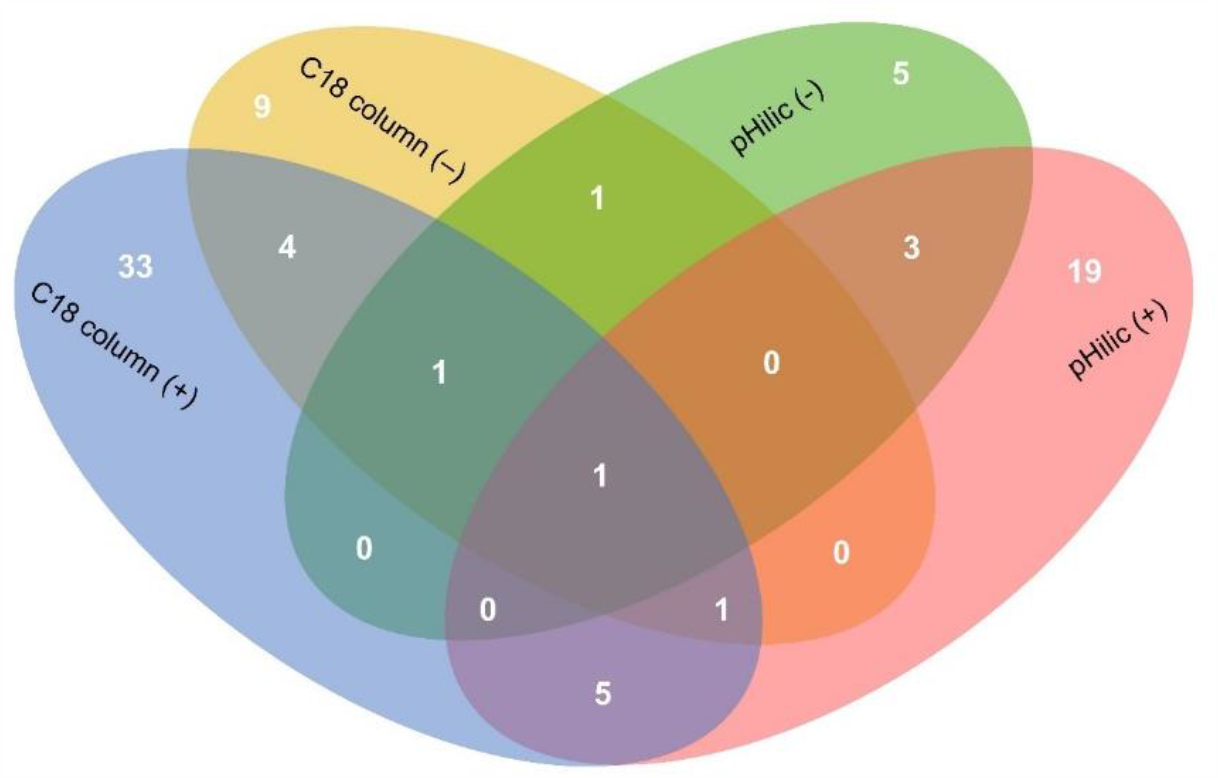
Venn diagram showing distribution of 82 metbolites in CSH-C18RP Column and ZIC-pHILIC column in both positive and negative ion modes.

Figure 4 shows example of the ion chromatograms of the selected metabolites and visualisation of their peaks in Compound Discoverer software. Images in the left column represent LC-MS peaks of the compounds (e.g., uric acid, D-glucose, L-pyroglutamic acid and urocanic acid) along with the adjusted retention times (RT). Images on the right column are the spectra of the respective compounds with their isotopes [M+H] ^+1^ and [M-H] ^-1^ for both column conditions. Regions highlighted in grey indicate a match to the theoretical isotope pattern.

**Figure 4.**
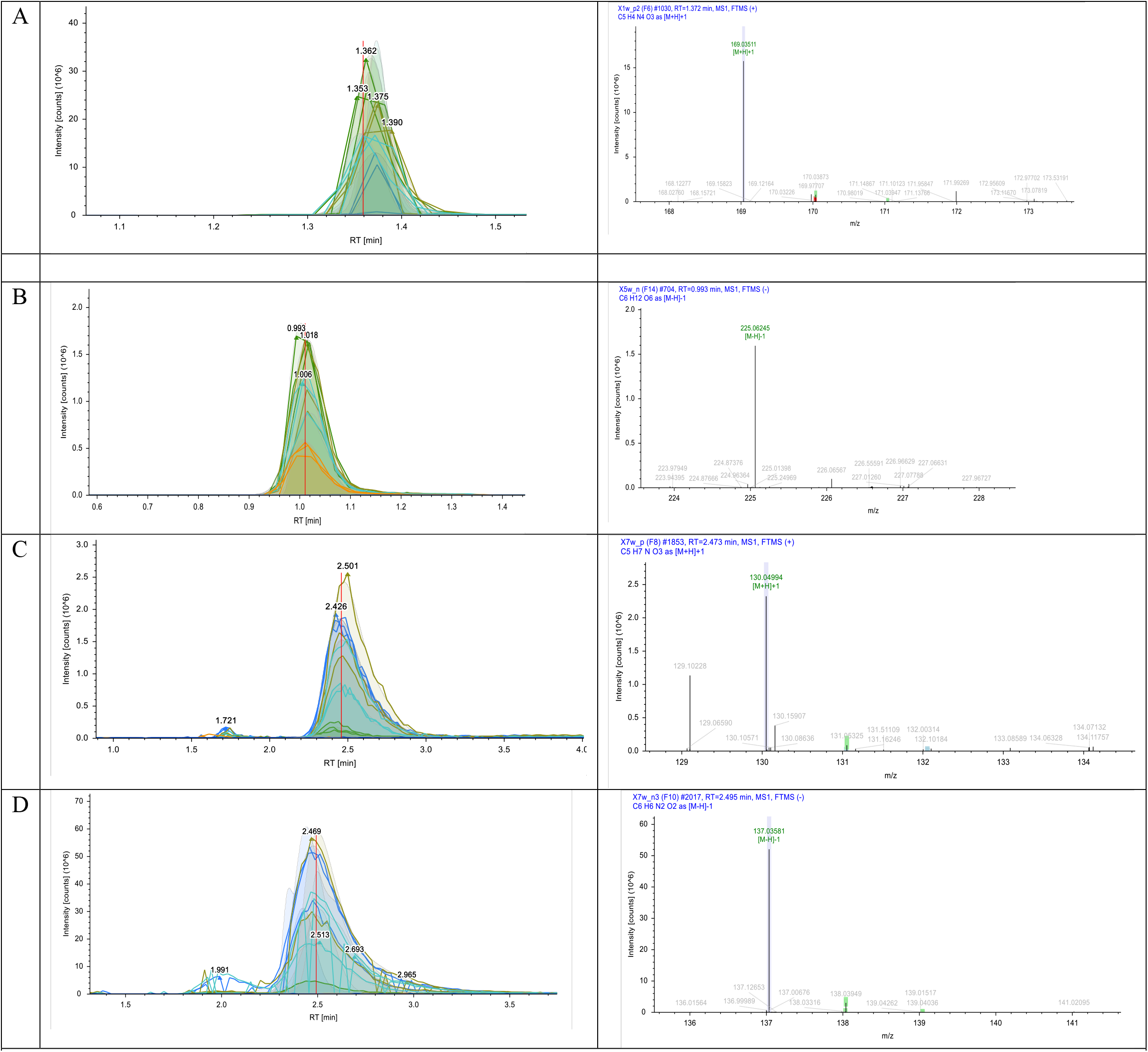
The output of an ion chromatogram on the Compound Discoverer. The left images represent mass spectrometry with an adjusted RT and the right image represents their isolated ion chromatograms. A) Uric acid on CSH-C18RP (+), RT = 1.372, m/z = 169.0351 [M+H]^+1^ B) D-Glucose on CSH-C18RP (-), RT = 0.993, m/z = 225.0624 [M-H]^-1^ C) L-Pyroglutamic acid on ZIC-pHILIC (+), RT = 2.473, m/z = 130.0499 [M+H]^+1^ D) Urocanic acid on ZIC-pHILIC (-), RT = 2.495, m/z = 137.0358 [M-H]^-1^

### Distribution of metabolites

Of 82 metabolites, 59 (71.5 %) were novel in the healthy tears (Supplement Table 2). Amino acids were the most frequently detected in the tears (23 / 82), which was followed by carbohydrates (10 / 82), carboxylic acids (7 / 82), carnitines (5 / 82) and glycerophospholipids (4 / 82). Amino acids, carboxylic acids, carnitines and glycerophospholipids were frequently detected in positive ion modes of both column conditions, whereas carbohydrates were detected mostly in negative ion modes of both column conditions. Moreover, 70 compounds (85.4 %) were present in all four samples with a CV of less than 25 %.

Similarly, figure 5 shows the relative abundance [in Log_2_ fold-change (FC)] of 82 tear metabolites. The average fold-change ± standard error was 5.4 ± 0.24 (Median = 5.3, 95% confidence level = 0.48). Tetrahydro-beta-carboline-3-carboxylic acid (Log_2_ FC = 12.72 ± 1.23, P_adj._ < 0.0001), hexitol (Log_2_ FC = 10.96 ± 0.46, P_adj._ < 0.0001) and 2-Hexenoylcarnitine (Log_2_ FC = 10.30 ± 1.89, P_adj._ < 0.0001) were the most abundant metabolites. Conversely, N-methyl ethanolamine phosphate (Log_2_ FC = 1.95± 0.21, P_adj._ <0.0001), D-glucose (Log_2_ FC = 2.2 ± 0.67, P_adj._ <0.0001) and 5-Chloro-4-oxo-L-norvaline (Log_2_ FC = 2.2 ± 0.72, P_adj._ <0.0001) were the least abundant metabolites.

**Figure 5:**
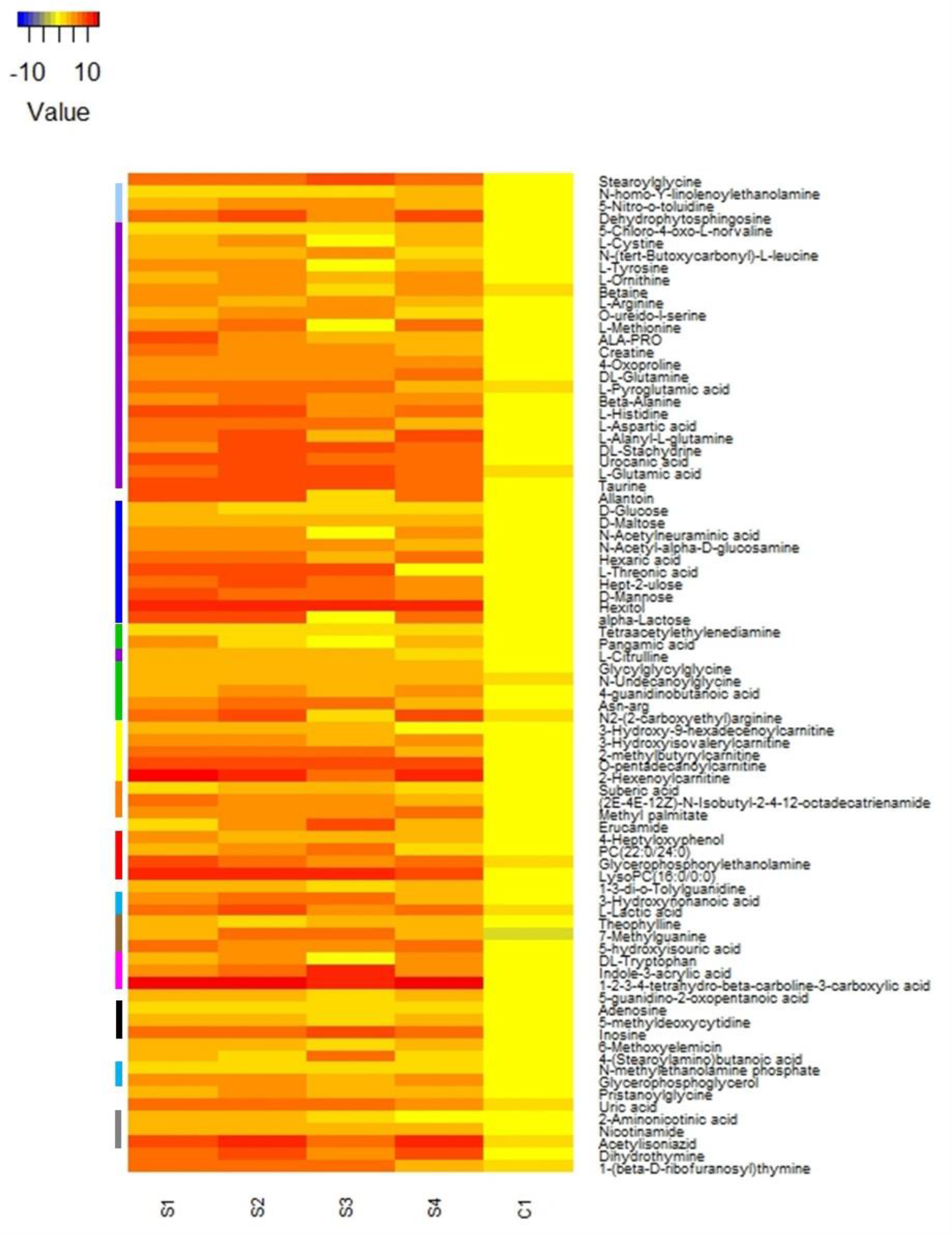
Relative abundance of 82 tear metabolites presented with Log_2_ fold-change at the CV less than 25% after B-H correction for false discovery rate at P < 0.05. Compound classes are represented with various colour codes in the left vertical column.

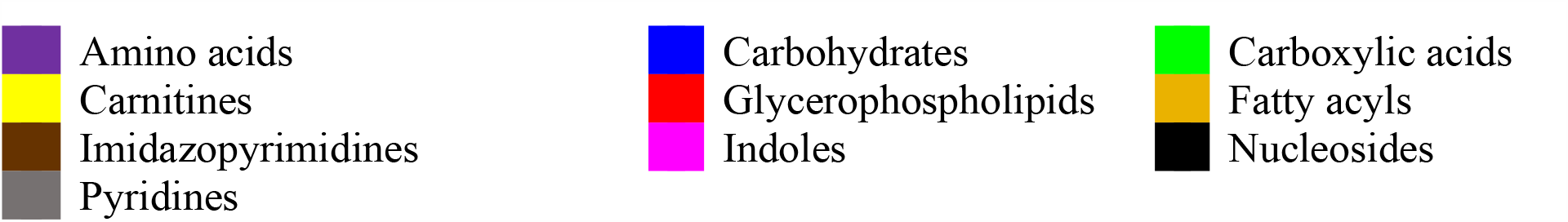

## Discussion

An untargeted metabolomics approach intends to comprehensively measure as many analytes as possible, including unknown chemicals. The current analytical method could cover a wide range of tear metabolome using complementary column chemistries (ZIC-pHILIC and CSH-C18RP) and ionisation (positive and negative), which resulted in 82 compounds classified into 23 compound classes.

Metabolite detection is dependent on the compound extraction method and data acquisition method and analytical method used. Chen et al. (2011) reported 60 metabolites from normal human tears collected with the Schirmer strip, using information-dependent acquisition (IDA) directed liquid chromatography-tandem mass spectrometry (LC-MS/MS) with a peak mining workflow built on Extracted Ion Chromatogram (XIC) manager (Chen et al., 2011). Similarly, Dammeier et al. (2018) quantified 104 metabolites from normal human tears collected with the Schirmer strip, using a metabolomics kit in the multiple reaction monitoring (MRM) platform (Dammeier et al., 2018). The present study acquired tandem mass spectrometric data (MS/MS) from automatic Data Dependant Analysis in Liquid chromatography-QExactive-HF hybrid quadrupole-Orbitrap-mass spectrometry (QE-HF-MS), and the data were analysed in Compound Discoverer software (Supplement Table 1). Of 82 metabolites, 59 metabolites (71.9 %) were novel in healthy tear fluid (Supplementary Table 1). The majority of novel metabolites were carnitines (4 out of 5 compounds), carbohydrates (9 out of 10 compounds) and carboxylic acids (7 out of 7 compounds).

CSH technology in C18RP incorporates a low-level surface charge to improve sample loading capacity, stability over a wide range of pH and superior peak shape for basic compounds. Moreover, acidic mobile phase in C18RP could elute around 90 non-polar compounds in positive ion mode using Fusion-RP column (Lu et al., 2006). In the present study, after filtering of data, 45 compounds were observed using the CSH-C18RP column in positive ion mode, including non-polar compounds such as DL-tryptophan, 2-hexenoylcarnitine, uric acid and L-tyrosine, and 12 compounds also detected in other ion modes (Figure 3). The difference in compound detection could be due to the analytical difference related to the application of the LC condition, elution gradient and the column chemistry. Previously, reversed-phase (RP) high-performance liquid chromatography (HPLC) with electrospray ionization (ESI) tandem mass spectrometry was used to profile and quantify the relative composition and concentrations of 23 amino acids from both basal and reflex tears (Nakatsukasa et al., 2011). In that study, tear fluid was analysed in multiple-reaction monitoring (MRM). The present study eluted 15 amino acids in CSH-C18RP column of HPLC with HESI, and overall 23 amino acids were eluted in the present analysis. Moreover, O-ureido-l-serine, N-undecanoylglycine, ala-pro, betaine, DL-stachydrine, L-alanyl-L-glutamine, N-(tert-Butoxycarbonyl)-L-leucine, asn-arg and L-aspartic acid were putatively identified as new metabolites in the tears in the present study.

Hydrophilic interaction chromatography (HILIC) is a valuable alternative to C18RP liquid-chromatography, to separate polar, weakly acidic or basic compounds (Grumbach et al., 2004; Jandera, 2011). The ZIC-pHILIC column utilises a zwitterionic stationary phase for separating highly polar hydrophilic compounds across a wide spectrum of pH. The present study eluted a wide range of metabolites such as amino acids, carbohydrates, carboxylic acids, carnitines, indoles and phosphate esters in ZIC-pHILIC similar to the findings in the previous studies (Lu et al., 2008). Therefore, the combination of both C18RP and pHILIC could complement the quantitation of a large group of both polar and no-polar compounds (Chen et al., 2011; Lu et al., 2008).

HESI enhances sensitivity in terms of ion counts and signal-to-noise ratio compared to regular ESI, by quickly evaporating initial droplets to expose the charged analyte of interest (Nguyen and Fenn, 2007). However, HESI performs poor for pyridine nucleotide from a cellular extract, probably due to heat-induced decomposition (Lu et al., 2008). In the present study, two pyridine compounds (acetylisoniazid and dihydrothymine) were detected.

## Limitations of the study

A single analytical method is not be sufficient to detect all metabolites in a biological sample. Combined analytical approaches are needed to maximise the number of metabolites observed. Therefore, metabolite detection in one study may vary with that of another one. Several factors are responsible for such variation such as strong adduct formation with inorganic ions such as sodium formate or potassium formate, the presence of a low concentration of certain metabolites, and difference in analytical approaches (Chen et al., 2011). The small sample size also confounds the generalisability of the study findings.

In conclusion, the current untargeted metabolomic platform is capable of detecting a wide range of human tear metabolites, including amino acids, carbohydrates, carboxylic acids, carnitines and glycerophospholipids. This approach may be applied for further metabolite studies of the ocular surface.

## Acknowledgement

The study was conducted under the Scientia Fellowship Program, the Scientia Graduate Research School, UNSW Sydney. Mass-spectrometric results were obtained at the Bioanalytical Mass Spectrometry Facility, UNSW, Sydney. Subsidised access to this facility is gratefully acknowledged.

## References

Ablamowicz, A.F., Nichols, J.J., 2017. Concentrations of MUC16 and MUC5AC using three tear collection methods. Mol. Vis. 23, 529.

Azkargorta, M., Soria, J., Acera, A., Iloro, I., Elortza, F., 2017. Human tear proteomics and peptidomics in ophthalmology: Toward the translation of proteomic biomarkers into clinical practice. J. Proteomics 150, 359–367.

Chen, L.Y., Zhou, L., Chan, E.C.Y., Neo, J., Beuerman, R.W., 2011. Characterization of the human tear metabolome by LC-MS/MS. J. Proteome Res. 10, 4876–4882.

ChenZhuo, L., Murube, J., Latorre, A., del Rio, R.M., 2000. Differential presence of amino acids in human tears of normal and dry eyes. Cornea 19, S80.

Choy, C.K.M., Cho, P., Chung, W.-Y., Benzie, I.F., 2001. Water-soluble antioxidants in human tears: effect of the collection method. Investigative Opthalmology and Visual Science 42, 3130–3134.

Dammeier, S., Martus, P., Klose, F., Seid, M., Bosch, D., Janina, D.A., Ziemssen, F., Dimopoulos, S., Ueffing, M., 2018. Combined Targeted Analysis of Metabolites and Proteins in Tear Fluid With Regard to Clinical Applications. Translational Vision Science Technology 7, 22–22.

Dor, M., Eperon, S., Lalive, P.H., Guex-Crosier, Y., Hamedani, M., Salvisberg, C., Turck, N., 2019. Investigation of the global protein content from healthy human tears. Exp. Eye Res. 179, 64–74.

Farkas, Á., Vámos, R., Bajor, T., Müllner, N., Lázár, Á., Hrabá, A., 2003. Utilization of lacrimal urea assay in the monitoring of hemodialysis: conditions, limitations and lacrimal arginase characterization. Exp. Eye Res. 76, 183–192.

Gogia, R., Richer, S.P., Rose, R.C., 1998. Tear fluid content of electrochemically active components including water soluble antioxidants. Curr. Eye Res. 17, 257–263.

Group, B.D.W., Atkinson Jr, A.J., Colburn, W.A., DeGruttola, V.G., DeMets, D.L., Downing, G.J., Hoth, D.F., Oates, J.A., Peck, C.C., Schooley, R.T., 2001. Biomarkers and surrogate endpoints: preferred definitions and conceptual framework. Clin. Pharmacol. Ther. 69, 89–95.

Grumbach, E.S., Wagrowski-Diehl, D.M., Mazzeo, J.R., Alden, B., Iraneta, P.C., 2004. Hydrophilic interaction chromatography using silica columns for the retention of polar analytes and enhanced ESI-MS sensitivity. Lc Gc North America 22, 1010-+.

Hagan, S., Martin, E., Enríquez-de-Salamanca, A., 2016. Tear fluid biomarkers in ocular and systemic disease: Potential use for predictive, preventive and personalised medicine. EPMA Journal 7, 15.

Hohenstein-Blaul, N.V.U., Funke, S., Grus, F.H., 2013. Tears as a source of biomarkers for ocular and systemic diseases. Exp. Eye Res. 117, 126–137.

Jandera, P., 2011. Stationary and mobile phases in hydrophilic interaction chromatography: a review. Anal. Chim. Acta 692, 1–25.

Lu, W., Bennett, B.D., Rabinowitz, J.D., 2008. Analytical strategies for LC–MS-based targeted metabolomics. Journal of Chromatography B 871, 236–242.

Lu, W.Y., Kimball, E., Rabinowitz, J.D., 2006. A high-performance liquid chromatography-tandem mass spectrometry method for quantitation of nitrogen-containing intracellular metabolites. J. Am. Soc. Mass Spectrom. 17, 37–50.

Nakatsukasa, M., Sotozono, C., Shimbo, K., Ono, N., Miyano, H., Okano, A., Hamuro, J., Kinoshita, S., 2011. Amino acid profiles in human tear fluids analyzed by high-performance liquid chromatography and electrospray ionization tandem mass spectrometry. Am. J. Ophthalmol. 151, 799-808. e791.

Nalbantoglu, S., 2019. Metabolomics: Basic Principles and Strategies, Mol. Med. IntechOpen.

Nguyen, S., Fenn, J.B., 2007. Gas-phase ions of solute species from charged droplets of solutions. Proc. Natl. Acad. Sci. U. S. A. 104, 1111–1117.

Pescosolido, N., Imperatrice, B., Koverech, A., Messano, M., 2009. L-carnitine and short chain ester in tears from patients with dry eye. Optom. Vis. Sci. 86, E132–E138.

Pieragostino, D., D’Alessandro, M., di Ioia, M., Di Ilio, C., Sacchetta, P., Del Boccio, P., 2015. Unraveling the molecular repertoire of tears as a source of biomarkers: beyond ocular diseases. Proteomics 9, 169–186.

Pintor, J., Carracedo, G., Alonso, C.M., Bautista, A., Peral, A., 2002. Presence of diadenosine polyphosphates in human tears. Pflugers Arch. 443, 432–436.

Pinu, F.R., Goldansaz, S.A., Jaine, J., 2019. Translational Metabolomics: Current Challenges and Future Opportunities. Metabolites 9.

Rantamaki, A.H., Seppanen-Laakso, T., Oresic, M., Jauhiainen, M., Holopainen, J.M., 2011. Human Tear Fluid Lipidome: From Composition to Function. PLoS One 6.

Roberts, L.D., Souza, A.L., Gerszten, R.E., Clish, C.B.J.C.p.i.m.b., 2012. Targeted Metabolomics. 98, 30.32. 31-30.32. 24.

Speek, A.J., van Agtmaal, E.J., Saowakontha, S., Schreurs, W.H., van Haeringen, N.J., 1986. Fluorometric determination of retinol in human tear fluid using high-performance liquid chromatography. Curr. Eye Res. 5, 841–845.

ThermoScientific, 2019. Compound discoverer user guide software version 3.1. Revision A XCALI-98120.

Van Haeringen, N., Glasius, E., 1977. Collection method dependant concentrations of some metabolites in human tear fluid, with special reference to glucose in hyperglycaemic conditions. Albrecht Von Graefes Arch. Klin. Exp. Ophthalmol. 202, 1–7.

Zhou, L., Zhao, S.Z., Koh, S.K., Chen, L., Vaz, C., Tanavde, V., Li, X.R., Beuerman, R.W., 2012. In-depth analysis of the human tear proteome. J. Proteomics 75, 3877–3885.

