## Supplementary table 1 for "Human Tear Metabolomics Using Liquid Chromatography-Q Exactive-HF Mass Spectrometry"

**Supplementary Table 2. A comprehensive list of tear metabolites identified in the present study**

| **Amino acids** | **Carboxylic acids** | **Keto acids** |
| --- | --- | --- |
| *Alanine ^A,B,C^ | ^§^Tetraacetylethylenediamine | ^§^5-guanidino-2-oxopentanoic acid |
| ^§^ALA-PRO | ^§^4-guanidinobutanoic acid |  |
| *Arginine ^A,B,C,D^ | ^§^Pangamic acid | **Nucleosides** |
| ^§^Betaine | ^§^N2-(2-carboxyethyl) arginine | *Inosine ^D^ |
| *Creatine ^D^ | ^§^Asn-arg | ^§^Adenosine |
| *Citrulline ^B,C,D^ | ^§^Glycylglycylglycine | ^§^5-methyldeoxycytidine |
| ^§^DL-Stachydine | ^§^N-Undecanoylglycine |  |
| *Glutamic acid ^A,B,C,D^ |  | **Phenol ester** |
| *Glutamine ^A,B,C,D^ | **Carnitines and Acylcarnitines** | ^§^6-Methoxyelemicin |
| *Histidine ^A,B,C^ | *Hexanoylcarnitine ^D^ |  |
| ^§^L-Alanyl-L-glutamine | ^§^O-pentadecanoylcarnitine | **Phenylpropanoic acid** |
| ^§^L-Aspartic acid | ^§^3-Hydroxyisovalerylcarnitine | ^§^4-(Stearoylamino)butanoic acid |
| *L-Cystine ^C^ | ^§^3-Hydroxy-9-hexadecenoylcarnitine |  |
| *Methionine ^B.C^ | ^§^2-methylbutyrylcarnitine | **Phosphate ester** |
| ^§^N-(tert-Butoxycarbonyl)-L-leucine |  | ^§^N-methyl ethanolamine phosphate |
| *Ornithine ^B.C^ | **Fatty Acyl** | ^§^Glycerophosphoglycerol |
| ^§^O-ureido-l-serine | ^§^Suberic acid |  |
| *Pyroglutamic acid ^D^ | ^§^Methyl palmitate | **Prenol** |
| *Taurine ^A,B,C,D^ | ^§^2,4,12-Octadecatrienoic acid isobutylamide | ^§^N-(2,6,10,14-Tetramethylpentadecanoyl) glycine |
| *Tyrosine ^B,C,D,E,F^ |  |  |
| *Urocanic acid ^D^ | **Fatty amide** | **Purines and derivatives** |
| ^§^4-Oxoproline | ^§^Erucamide | *Uric acid ^D,E,F,K^ |
| ^§^5-Chloro-4-oxo-L-norvaline |  |  |
|  | **Glycerophospholipid** | **Pyridine** |
| **Acylglycin** | * lysoPC 16:0 ^B^ | ^§^Acetylisoniazid |
| ^§^Stearoylglycine | ^§^4-Heptyloxyphenol | ^§^2-Aminonicotinic acid |
|  | ^§^PC (22:0/24:0) | ^§^Nicotinamide |
| **Amines** | ^§^Glycerophosphoryl ethanolamine |  |
| ^§^N-homo-Y-linolenoyl ethanolamine |  | **Pyrimidine** |
| ^§^5- Nitro-o-toluidine | **Guanidine** | ^§^Dihydrothymine |
| ^§^ (8E)-2-Amino-8-octadecene-1,3,4-triol | ^§^1,3-di-o-Tolylguanidine |  |
|  |  | **Pyrimidine nucleoside** |
| **Amino Ketones** | **Hydroxy acids** | ^§^1-(beta-D-ribofuranosyl) thymine |
| *Allantoin ^D^ | *Lactate/Lactic acid ^G^ |  |
|  | ^§^3-Hydroxynonanoic acid |  |
| **Carbohydrates** |  |  |
| ^§^N-Acetyl-alpha-D-glucosamine | **Imidazopyrimidine** |  |
| ^§^Hept-2-ulose | ^§^Theophylline |  |
| ^§^Hexaric acid | ^§^5-hydroxyisouric acid |  |
| ^§^D-Maltose | ^§^7-Methylguanine |  |
| ^§^Hexitol |  |  |
| ^§^D-Mannose | **Indoles** |  |
| ^§^L-Threonic acid | ^§^Indole-3-acrylic acid |  |
| ^§^α-Lactose | *Tryptophan (amino acid) ^B.C.D^ |  |
| *Glucose ^G,I,J^ | ^§^1,2,3,4-tetrahydro-beta-carboline-3-carboxylic acid |  |
| *N-Acetylneuraminic acid ^D^ |  |  |
| * Also reported in the literature  ^§^ First time found in the present study  A = ChenZhuo et al., 2000; B = Dammeier et al., 2018; C = Nakatsukasa et al., 2011; D = Chen et al., 2011; E = Choy et al., 2001; F = Gogia et al., 1998; G = Van Haeringen and Glasius, 1977; H = Pescosolido et al., 2009; I = Baca et al., 2007; J = Taormina et al., 2007; K = Mendelsohn et al., 1998 | | |
